# JiSuJi, a virtual muscle for small animal simulations, accurately predicts force from naturalistic spike trains

**DOI:** 10.1101/2025.06.16.659528

**Authors:** Weili Jiang, Iris Adam, Nicholas W. Gladman, Sam Sober, Qian Xue, Coen P.H. Elemans, Xudong Zheng

**Author notes:** Department of Musculoskeletal and Ageing Science, Institute of Life Course and Medical Sciences, University of Liverpool, Liverpool, United Kingdom (Nicholas W. Gladman).

## Abstract

Physics-based simulators for neuromechanical control of virtual animals have the potential to significantly enhance our understanding of intricate structure-function relationships in neuromuscular systems, their neural activity and motor control. However, a key challenge is the accurate prediction of the forces that muscle fibers produce based on their complex patterns of electrical activity (“spike trains”) while preserving model simplicity for broader applicability. In this study, we present a chemomechanical, three-dimensional finite-element muscle model – JiSuJi (pronounced *jì sù jī*, meaning “ultrafast muscle” in Chinese) - that efficiently and accurately predicts muscle forces from naturalistic spike trains. The model’s performance is validated against songbird vocal muscles, a particularly fast and therefore challenging muscle type. Our results demonstrate that JiSuJi accurately predicts both isometric and non-isometric muscle forces across a variety of naturalistic neural activity patterns. JiSuJi furthermore outperforms state-of-the-art muscle simulators for accuracy, while maintaining computational efficiency. Simulating muscle behavior offers a promising approach for investigating the underlying mechanisms of neuro-muscular interactions and precise motor control, especially in the fast-contracting muscles of animal model systems.

## Introduction

Animals precisely control their movements to execute a wide array of behaviors, but the neuromuscular mechanisms, which describes how the nervous system controls muscle activity to generate movement, are still not well understood. Unraveling these mechanisms is crucial for advancing our understanding of motor control, diagnosing movement disorders, and developing effective bioinspired systems and rehabilitation technologies. Recent advances in simulation techniques opened the exciting possibility to build physics-based simulators for neuromuscular control of virtual bodies (1–5) that are grounded in anatomically and biomechanically accurate models, including whole-body simulations of drosophila (6), mouse (7) and rat (8). In parallel, rapid advances in AI-driven, three-dimensional (3D) posture extraction of animal bodies, high-density electromyography and neural recording allows unprecedented large-scale datasets of simultaneous body kinematics, muscle and brain activity to validate prediction of such embodied models (9–16). Together, these tools will be instrumental in increasing our understanding of complex structure-function relationships within neuromuscular systems, predictions of neural activity, and motor control principles.

Key to accurate predictions of body posture and behavior is the amount of force produced by virtual muscles. However, current muscle models are not able to use natural, single-unit electromyography (SU-EMG) or neural data as inputs (17,18). A commonly used muscle force model is the Hill-based muscle activation model (19,20) that predicts muscle force using a single input variable representing the overall activation level of a muscle, typically estimated *via* bulk electromyography (17). Its simplicity and low computational demands allow quick force approximations in 1D string (2,5,21) and in 3D finite-element implementations (18,22), but the input of this model approach does not allow variable interspike intervals (ISIs) and non-tetanic stimulation, as found in natural spike trains (23). In contrast, more complex physics-based models (24–26) incorporate the electrochemical processes underlying neural signaling, as described by the classic Hodgkin-Huxley model (27), and link them with skeletal muscle contraction, providing detailed representations of the excitation-contraction coupling (ECC) pathway from motor endplate activation to force generation in muscle. These models can take variable ISI’s and non-tetanic stimulation as input (24–26), but their complexity makes them computationally intensive. Furthermore, these models require parametrization of numerous parameters, many of which have only been measured for specific muscles (22,28). These constraints limit their broad practical applications in complex 3D musculoskeletal simulations. We thus lack muscle force models that allow naturalistic neural input while also balancing computationally efficiency and force prediction precision.

Vertebrate muscles consist of a mosaic of muscle fiber types. Simulating force generation of particularly fast twitch muscles, which are prevalent in muscles specialized for rapid movements (29) in vertebrate animal models such as mouse, rat and songbirds, remains challenging. Driven by recent developments in high-density EMG arrays capable of recording single-unit muscle activity, there is growing evidence that spike timing in motor units (the muscle fibers innervated by a single motor neuron) plays a significant role in controlling motor behavior across various species, and particularly so in fast muscles (30–33). This further emphasizes the need that next generation muscle simulators can take naturalistic spike trains as input.

Spike trains recorded from muscle fibers during behavior have revealed that the precise statistics of spiking strongly modulates muscle force and the resulting behavior, including millisecond-precise regulation of ISIs (30,31,34–36)). Some muscle models can use spike trains with variable ISI’s as input to predict muscle force (37–45). However, these models often rely on simplified representations of the underlying excitation-contraction coupling (ECC) processes, using submodels that differ in how they approximate neural input, calcium handling, troponin activation, and force generation. Consequently, some models exhibit reduced accuracy in predicting force responses to multiple-pulse spike trains (40,41) or low- to mid-frequency spike trains (42). To our best knowledge, the only current muscle model that can use long, irregular spike trains, different muscle lengths as input and generated a force response that matched experiment measurement, requires quantification of over thirty parameters (43) that are not commonly available. We thus lack a muscle model that can emulate the fast twitch muscles of common laboratory animal models.

Here, we present a novel chemomechanical muscle force model that overcomes this challenge and allows for accurate and efficient prediction of muscle forces from naturalistic spike trains. To extend its application to muscle contractions during length changes, which is essential for modeling virtual bodies in action, we integrated this muscle force model into a 3D finite element framework (46,47) that is able to incorporate the change of muscle length. We refer to this chemomechanical, 3D finite-element muscle model as JiSuJi (极速肌), Chinese for Fast Muscle Simulator. We validated the model performance in syringeal muscles of songbirds, known for their millisecond precise neuromuscular control (48,49), and therefore particularly challenging. For isometric contractions our model outperforms state-of-the-art models and accurately predicts muscle force patterns under diverse spike patterns, including those with constant and variable ISIs across both short (1-3 spikes) and long durations. For non-isometric contractions, we validated a 3D finite element implementation of the model against experimental muscle where we precisely quantified the geometry using an ultra-high-frequency ultrasound scanner. Our model also accurately predicted non-isometric performance of a 3D muscle. The model can be trained on parameters that are easily obtained from experimental isometric and isovelocity force-length data. Together, these features make our model a promising tool for studying the mechanisms underlying complex neuro-muscle interactions in precise motor control.

## Results

### Muscle model approach

We developed a muscle model that includes the four critical processes of the ECC pathway underlying muscle force production: (1) the change of calcium channel membrane permeability upon action potential (AP) arrival at the neuromuscular junction and sarcolemma (the muscle fiber membrane); (2) the resultant change in free calcium concentration [Ca^2+^] within the sarcoplasm (the muscle cell cytoplasm); (3) the binding and unbinding of calcium to troponin, whose calcium-induced conformation change frees up myosin binding sites on actin; and (4) force generation through actin-myosin cross-bridge cycling, where the actin–myosin interaction enables the actin to slide over the myosin and tension production (50) (See **Methods**). During rapid successive APs, calcium release from the sarcoplasmic reticulum (SR), the membrane-bound structure within muscle cells that stores [Ca^2+^], is larger than active calcium reuptake or buffering mechanisms, which increases the intracellular free [Ca^2+^] and thereby force (51,52). Once AP firing rate exceeds a certain threshold, [Ca^2+^] reaches a maximum value that allows all available binding sites to form cross-bridges and the muscle will produce its muscle-specific maximal isometric force, also known as tetanic force. Thus accurately predicting force dynamics from naturalistic AP inputs – from sparse irregular spikes to tetanic maximal stimulation – crucially requires the inclusion of intracellular [Ca^2+^].

Furthermore, we introduce a novel component that accounts for dynamic calcium release from the SR due to different ISIs building on previous work (40). We implemented a genetic algorithm-based optimization approach that estimates the model parameters by minimizing the difference between measurement and prediction in a small subset of experimental stimulation patterns (see **Methods**).

### Isometric force predictions

We tested our model’s predictive performance in songbird vocal muscles that control airflow and tension of vibratory tissue (53,54) to control sound production in the avian vocal organ. These muscles are superfast muscles that can resolve millisecond-scale variations in spike timing to regulate respiration and song (48,49). Their superfast calcium dynamics and force development (55) require an accurate prediction of muscle force generation under varied ISIs (33–35), which is a particular challenge for current models.

First, we tested our model’s predictive accuracy for isometric force production in response to short duration stimulation with varied ISIs. As experimental benchmark we used previously published isometric force dataset from isolated muscle fiber bundles of the dorsal tracheobronchial (DTB) muscle from the zebra finch (33). Muscle fiber bundles were subjected to a large array of twenty-five three-spike stimulation patterns with varied ISIs. Based on (35), three-spike stimulation patterns were chosen as the minimal set that maintain fixed firing rate, burst onset, and burst duration, while allowing variation in spike timing through manipulation of the middle spike’s timing. Our model simulations accurately predicted the time-course of isometric force development in experimentally measured responses (**Fig. 1a**). The high prediction accuracy of the model becomes apparent when comparing across spike patterns. Linear regressions between three predicted and measured statistics, i.e., peak force (F_max_), area under the curve (AUC), and time to reach peak force (t_p_) have slope *k* =1.00, 1.00, and 0.99 and R^2^ = 0.96, 0.94 and 1.00, respectively, across all spike patterns (**Fig. 1b**). F_max_, AUC and t_p_ did not differ significantly between predicted and measured values (Paired t-tests, F_max_: p=0.86, AUC: p=0.73; t_p_: p=0.07, n=24 patterns). The variability of predicted force production is on par with or even less than the variability between technical replicates of spike patterns in the experimental data (48). Our model thus accurately predicts key parameters describing time-resolved force development for short duration stimulations with varied ISIs.

**Figure 1.**
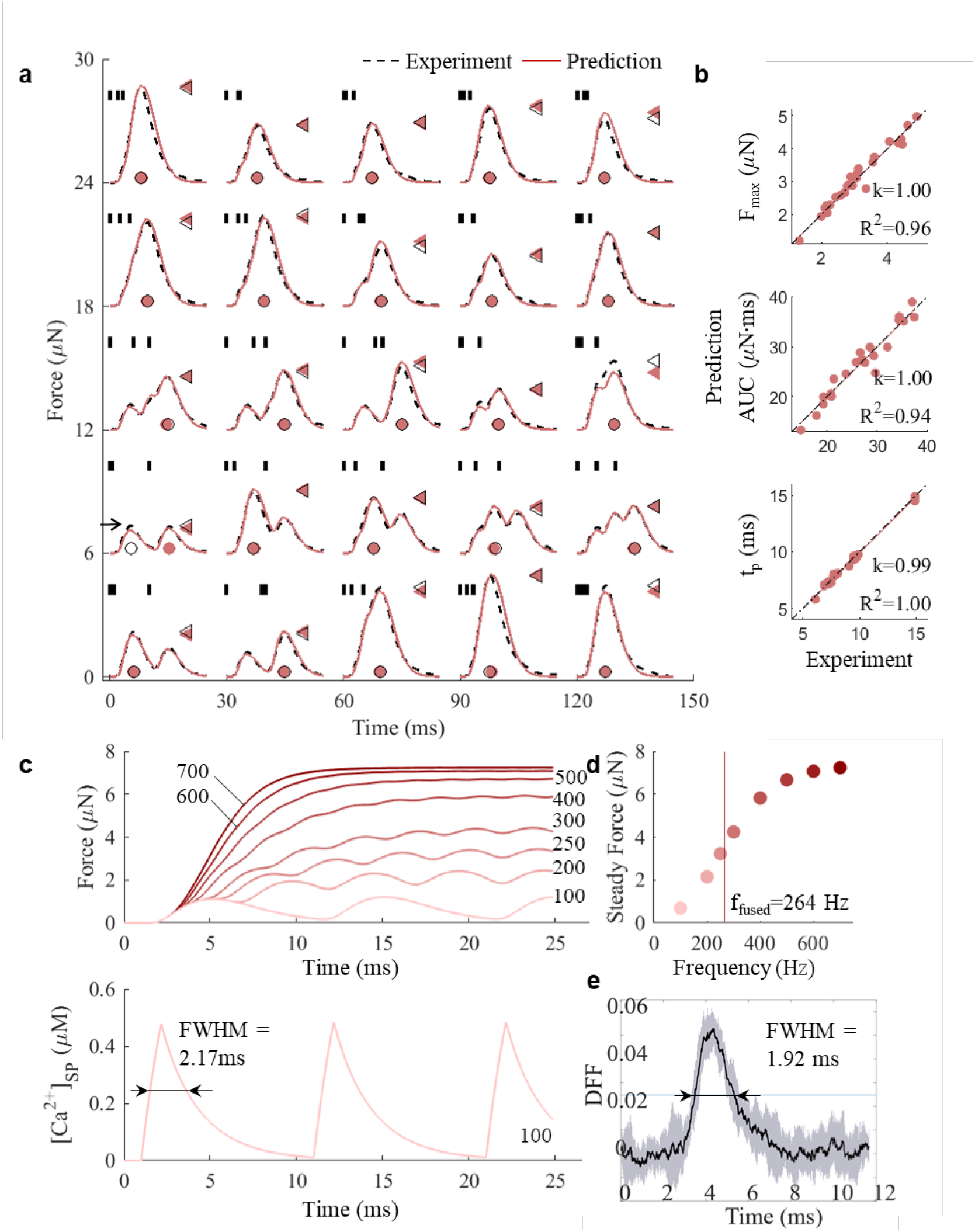
JiSuJi accurately predicts isometric force production. **a**, Comparison of forces generated in response to twenty-five three-spike stimulation patterns (vertical short bars) in the zebra finch dorsal tracheobronchial (DTB) muscle from (33). Maximum force (F_max_, triangles) is reached at time t_p_ (circles). **b**, Simulations accurately predict experimental data for F_max_, area under curve (AUC) and t_p_ for stimulation patterns in panel **a**. One case was excluded (horizontal arrow) because peak force values correspond to different incoming spikes. **c**, Long duration stimulation with constant ISIs – force (top) and intracellular calcium concentration corresponding to 100Hz stimulation (bottom). FWHM: full width at half maximum. **d**, Tetanic force increases with stimulation frequency and fuses when fusion index (FI) equals 0.9 at 264 Hz. **e**, Relative intracellular [Ca^2+^] signal (mean (black line) ± std (gray area) of 10 overlaid twitches of 100 Hz stimulation in one individual with FWHM = 1.92 ± 0.17 ms at 39°C (55) consistent with simulation in **d**.

Next, we tested our model’s predictive accuracy for isometric force production in response to long duration stimulations with constant ISIs. Long duration spike trains have been particularly challenging for previous models (40–42) that suffered from limited accuracy in dynamic force prediction to multiple-pulse spike trains (40,41) and at low- to mid-frequency spike trains (42). Our simulations show that as AP firing rate increases over 100 Hz, the force responses from individual twitches-muscle force produced by a single neural spike, starts to fuse, which results in incomplete relaxation between contractions (**Fig. 1c**; **Table S1**). The maximum force increases with the stimulation frequency between 100 and 700Hz, with the rate of increase notably slowing beyond 400Hz (**Fig. 1d**), consistent with previous observations (36,49). To compare the progression of tetanic fusion to experimental observations, we calculated the fusion index (FI), which is defined as the ratio of minimum to maximum force produced between two consecutive stimuli (56). The predicted fusion frequency (f_fused_ at FI =0.9) is 264 Hz, which falls within the experimentally observed range of 200-300 Hz (33). Two key features of tetanic contraction-deceleration of force increase as it nears tetanic contraction and f_fused_ thus align well with experimental observations (33).

Simulated intracellular calcium [Ca^2+^] traces show fully separated transients at 100 Hz stimulation (**Fig. 1c**) with a full width at half maximum (FWHM) value of 2.17 ms of individual twitches. Because muscle shortening induces severe motion artifacts in [Ca^2+^] measurements, [Ca^2+^] transient dynamics can only be measured accurately when force production is prevented using actomyosin interaction inhibition (57). It is therefore not possible to measure force production and [Ca^2+^] simultaneously. We previously measured [Ca^2+^] using high-affinity calcium dye in zebra finch DTB muscle (55). During stimulation series at 100Hz, the [Ca^2+^] were fully separated and resulted in [Ca^2+^] FWHM of 1.98 ± 0.65 ms (N=4 individuals) at 39°C (**Fig. 1e**, See **Methods**). Our predicted values from fitting force traces fit well to these experimental observations. Taken together, our muscle model accurately predicts isometric force production for both short duration stimulations with varied ISI and fusion properties of long duration stimulations with constant ISIs.

The previous benchmark datasets were, however, not collected in the same individuals, which prevents direct comparison of time-resolved tetanic properties. To quantitatively compare the predictive capacity of our -and previous models- for time-resolved isometric force development in both short and long duration stimulations, we collected a new experimental benchmark dataset that contains both paradigms (long duration stimulations with constant ISIs and short duration stimulations with varied ISIs) in the same individuals. To test if our model generalizes across muscles and across species, we collected data in another vocal muscle, the ventral syrinx (VS) muscle, in another songbird species, the Bengalese finch. Again, we tested our model’s predictive accuracy for isometric force production in response to short duration stimulations with varied ISIs.

Our model simulations accurately predict the time-course of isometric force development across ten short duration stimulations (up to three spikes) with varied ISIs (**Fig. 2a; Table S1**). The three key predicted statistics F_max_, AUC, and t_p_ describing the force curves were not significantly different between simulation and experiment across the spike patterns (Paired t-tests, F_max_: p=0.35; AUC: p=0.11; and t_p_: p=0.21, n=9 patterns) and had strong correlation between predicted and measured values (linear regressions: slope *k* =1.03, 1.04, and 0.99; R^2^ = 0.95, 0.98 and 0.99 for F_max_, AUC, and t_p_, respectively) across all spike patterns (**Fig. 2b**).

**Figure 2.**
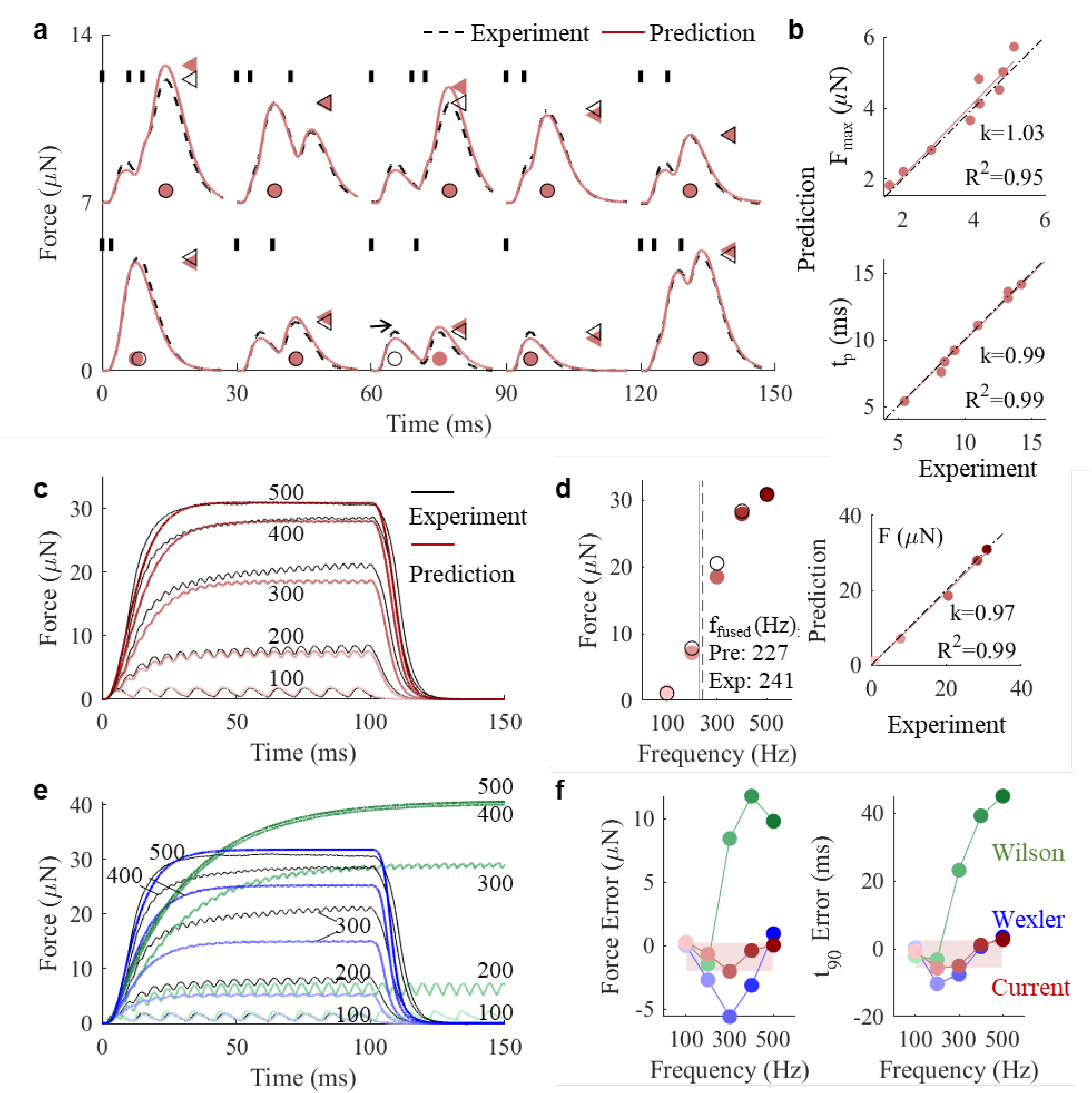
JiSuJi outperforms state-of-the-art models on force production by naturalistic input. **a**, Comparison of forces generated in response to ten short duration stimulation patterns (vertical short bars) in the Bengalese finch ventral syringeal (VS) muscle. Maximum force (F_max_, triangles) is reached at time t_p_ (circles). One case was excluded (horizontal arrow) because peak force values correspond to different incoming spikes. **b**, Simulations accurately predict experimental data for F_max_ and t_p_ for stimulation patterns in panel **a. c**, Time traces of simulated and measured isometric forces in response to long duration stimulations with constant ISIs at different stimulation frequencies. **d**, Steady-state forces increase with stimulation frequency (left), and simulations accurately predict experimental data for steady force (right). **e**, Isometric force predictions by Wexler (blue) and Wilson (green) models. **f**, Force error between prediction and benchmark experimental dataset shows that our model outperforms two state-of-the-art models for steady force (left) and t_90_ (right).

For long duration stimulations with constant ISIs, the model accurately predicted the steady force magnitude achieved per frequency setting (linear regression: slope *k*=0.97; R^2^=0.99; **Fig. 2cd**). The predicted f_fused_ is 227 Hz is well-aligned with the experimental value of 241 Hz (**Fig. 2d**) and force error remains under 2μN (7% of tetanic force) across all frequency settings (**Fig. 2f**). The time error required to reach 90% steady force (t_90_) is predicted to occur within 6 ms (**Fig. 2f**).

These results confirm that our muscle model accurately predicts isometric muscle force generation across diverse stimulation patterns, ranging from constant and varied ISIs as well as short and long duration stimulations. By successfully applying the model to two distinct muscles from different species, we demonstrate the model’s capacity to generalize across both muscle types and species.

### Comparison to state-of-the-art muscle models

A widely used class of phenomenological muscle models, knows as activation dynamics models, represent muscle activation using first-order ordinary differential equations (ODEs) (20,58–60) and have been integrated into popular software – such as OpenSim (2) and Mujoco (5). The input to these models is typically a continuous excitation signal bounded between 0 and 1, often estimated from bulk electromyography recordings. However, because these models cannot simulate dynamic force responses to action potentials, we cannot directly compare their performance against JiSuJi. While a few studies (61– 66) have attempted to incorporate spike trains into this class of models, they failed to accurately reproduce force time courses at low stimulation rates (61).

After carefully reviewing the literature (17,18,22,61), we selected two representative state-of-the-art muscle models that accept spike train input, the Wilson model (42) and the Wexler model (40), to benchmark the performance of JiSuJi. The Wilson model was developed to describe the isometric force response of the locust hind leg extensor tibia muscle to fast extensor tibia motor neuron stimulation (67), and could describe the response of both fast and slow motor neuron stimulation (42,61). It employs a phenomenological first order ODE to capture the response of calcium concentration to spike train input and a sigmoid function to capture the nonlinear saturation between calcium concentration and the force response (17). This simplified, phenomenological approach contrasts with JiSuJi, which derives its governing equations directly from physiological process.

The Wexler model was developed based on rat skeletal muscle and demonstrated in soleus muscle and gastrocnemius muscle, which are slow and fast muscles, respectively. We selected the Wexler model (40) because it *i)* captures physiological processes underlying ECC and thus *ii)* can take naturalistic spike trains with variable ISI’s as input, and *iii)* its parameter type and number is feasible to determine experimentally. Although there are more recent implementation of the Wexler model (45,68), the basic model approach remains unchanged and we chose to follow the original model here. Other models were deemed either overly simplified (20), lacking biochemical process modeling (37–39), or too complex with too many unknown (e.g. over 30) parameters (43) to determine experimentally.

We fitted the Wilson model and the Wexler model to our experimental data using the same approach with JiSuJi, including the same dataset for fitting, objective function and genetic algorithm optimization method (See **Methods; Table S2**). The results from the Wilson model shows that for short duration stimulation patterns (**Fig. S1a**), there was no significant difference between simulation and experiment across stimulation patterns regarding F_max_ and AUC (Paired t-tests, F_max_: p=0.27; AUC: p=0.93; n=10 patterns). It should be noted that during relaxation, the predicted force did not follow the experimental data well. Force decrease after the last AP was too steep initially, followed by a too long tail compared to experimental data (**Fig. S1a**). However, when activated with long stimulation trains this model poorly predicts force development. The steady force magnitude error is up to 33% of tetanic force and 6 times larger than JiSuJi. The *t*_*90*_ occurs at 67 ms, long after the experimental data has reached steady force (**Fig. 2ef**), and 8 times poorer than JiSuJi.

The results from the Wexler model shows that for short duration stimulation patterns (**Fig. S1b**), F_max_ is correctly predicted and not significantly different from experiment data across stimulation patterns (Paired t-tests, p=0.17, n=10 patterns). However, AUC predictions are significantly different from experiment data (Paired t-tests, p=0.002, n=10 patterns). During long trains, error of force and *t*_*90*_ from the Wexler model exhibited V-shaped patterns – largest at 200/300Hz and smallest at 100 and 500Hz (**Fig. 2ef**). At 300Hz, force error from Wexler model was 2.8 times that of JiSuJi (**Fig. 2f**).

Taken together, optimized with the same technique and validated against the same experiment dataset with both short and long duration stimulation patterns, JiSuJi outperforms these state-of-the-art models in predicting the force responses to short duration stimulations, and demonstrates markedly superior performance in predicting long duration stimulations with more precise steady force magnitude and timing.

### 3D non-isometric force predictions

We next tested if JiSuJi can predict the force response during more complicated non-isometric muscle contractions where muscle length is changing. To simulate non-isometric contractions, we integrated our muscle force model into a 3D finite element framework (46,47) (**See Methods**) and validated this 3D model implementation against experimental data.

First, we measured the passive and active stress-strain relationship, and stress-velocity relationship of an isolated muscle fiber bundle using the isovelocity shortening paradigm (**Fig. S2; Table S3**), where muscles are exposed to a series of shortening ramps at constant velocity during stimulation (48) (**See Methods**). The 3D simulation requires an accurate 3D geometrical representation of the muscle bundle tested. Therefore, we used high-resolution 71 kHz ultrasound imaging (**See Methods**) to nondestructively quantify the 3D geometry of the live muscle fiber bundle after the experiment (**Fig. 3a**). This imaging technique does not require tissue fixation and therefore does not damage or shrink the muscle tissue. With the resulting structural scan we next constructed a finite-element mesh of the tested muscle bundle (**Fig. 3a**). A previously developed force–length model and force–velocity model (46,69) were adopted, and their parameters were separately optimized to match measurements (See **Methods**; **Fig. S2; Table S3**).

**Figure 3.**
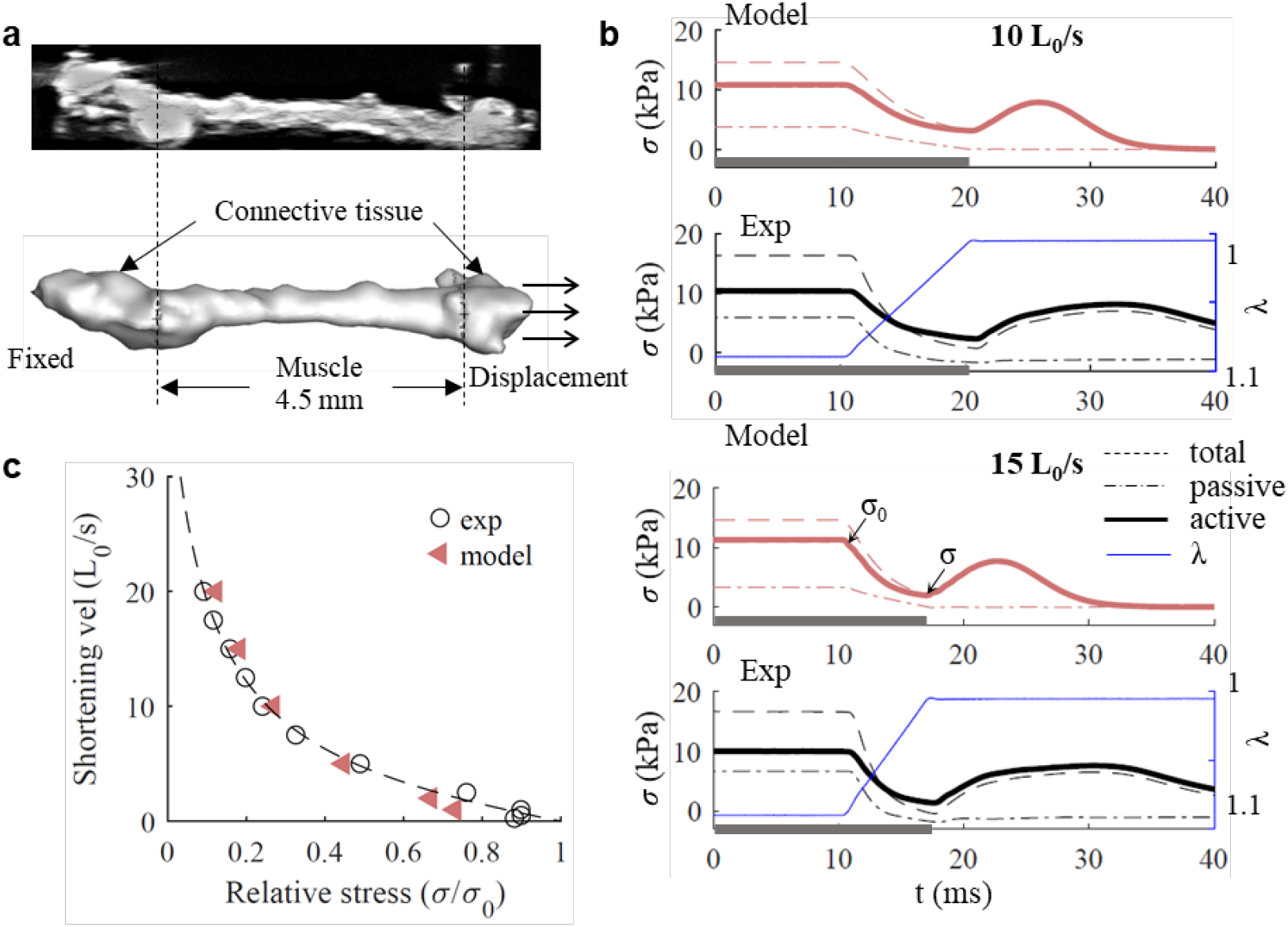
JiSuJi accurately predicts non-isometric force production. **a**, High-resolution 71MHz ultrasound scan sections (top) were used to quantify the 3D geometry of the living muscle preparation (bottom). Connective tissue at either end were boundary conditions in the simulation. **b**, Simulated and experimental total (dashed line), passive (dash dotted line) and active (solid line) stress during length changes (blue) at two representative shortening velocities, 10 L_0_/s and 15 L_0_/s. In both simulation and experiment, the muscle was stimulated at 600Hz. The time duration with the muscle stimulated was denoted using a horizontal grey bar along the x axis. **c**, The relative active stress (σ/σ_0_) – shortening velocity relationship from predictions and measurements. The dashed line described the experiment results by fitting a hyperbolic-linear equation (48). σ/σ_0_ was calculated using the active stress at the end of the shortening period (σ) divided by the active stress right before the shortening (σ_0_).

Next, we simulated isovelocity shortening of a zebra finch syrinx muscle bundle. JiSuJi accurately predicts active and passive stress as a function of time during isovelocity shortening (**Fig 3b**). Only after the shortening protocol the predicted active stress drops quicker in JiSuJi than in the experimental data (**Fig 3b**). Also the force-velocity curve is well-captured (**Fig. 3c**) and there was no significant difference between simulation and experiment (Paired t-tests, p=0.36, n=6 conditions).

In summary, our model accurately predicted the non-isometric muscle active stresses of the zebra finch DTB muscle during isovelocity shortening by employing a finite element model with realistic geometry. The model successfully captured the dynamic stress responses related to the shortening period in various shortening velocity conditions. Our model thus, for the first time offers the opportunity to build more comprehensive and realistic models to simulate complex motor control.

## Discussion

We present JiSuJi, a virtual muscle simulator capable of accurately and efficiently predicting force generation from naturalistic spike trains. The model’s performance was validated in fast-contracting muscles, syringeal muscles of songbirds, which exhibit millisecond precise neuromuscular control (48,49). To demonstrate its generalizability across muscle types and species, we applied JiSuJi to two distinct muscles from different songbird species. The model can be trained on parameters that are easily obtained from experimental isometric and isovelocity force-length data. Our model thus offers the opportunity to build comprehensive and realistic muscle models as a promising tool for simulating complex motor control.

Compared to state-of-the-art models (40,42), JiSuJi demonstrates superior accuracy in predicting isometric force responses across a range of spike patterns, including both constant and variable ISIs over short and long durations. We attribute this improvement primarily on the model’s incorporation of calcium-dependent inactivation (CDI) within calcium release dynamics. Calcium release can be reduced by elevated cytoplasmic calcium levels through CDI (70,71) or during high action potential spike rates, i.e. short ISI’s (72–74). Unlike previous models, which either assumed a constant SR permeability upon channel opening or relied on nonlinear relationships between calcium concentration and force generation to account for tetanic force saturation (38,42,43,75), JiSuJi dynamically adjusts calcium release based on ISI. This adjustment substantially enhances the model’s accuracy over previous models (40,42) in predicting force responses during prolonged stimulation. Importantly, our approach preserves key intermediate parameters, such as calcium concentration and activated troponin levels, and thus adds flexibility for further experimental fine-tuning.

The integration of JiSuJi into a 3D finite element framework enables accurate predictions of the dynamic stress responses related to the shortening period in various shortening velocity conditions of a muscle with accurate 3D geometry. The time courses of the active stress during isovelocity shortening are well-captured. Notably, while JiSuJi accurately predicts many aspects of active stress during shortening, minor deviations from experimental data were observed post-shortening. Specifically, the simulated active stress decreases due to reduced intracellular calcium levels ([Ca^2+^]), whereas experimental data indicate a prolonged high level of active stress before a gradual decline. This discrepancy may stem from history-dependent force development. Experimental findings suggest that, when stimulation is maintained tetanically after an isovelocity ramp, the degree of active force recovery following shortening is velocity-dependent (48). This implies that cross-bridge state dynamics may be influenced by shortening velocity. History-dependent force modulation could be associated with factors such as lattice spacing, thick filament stiffness, or non-cross-bridge forces (76,77), though the precise mechanisms remain unknown.

By incorporating JiSuJi into a 3D finite element framework, our model enables the simulation of non-isometric contractions in a realistic 3D setting—an essential feature for musculoskeletal modeling in virtual animals (21,78). Current physics-based simulators, including MuJoCo (5) and OpenSim (2), predominantly rely on Hill-type muscle models, which predict force production based on bulk muscle activation but do not account for neuromechanical processes such as transient calcium dynamics, length- and velocity-dependent calcium signaling, or motor unit recruitment (17). As a result, these simulators are limited to representing general activation levels rather than naturalistic spike timing. JiSuJi overcomes these limitations by incorporating precise spike timing in a computationally efficient manner. Additionally, it accounts for anatomical muscle properties, such as fiber density, and can be extended to include more realistic motor unit distributions (79–81). By bridging the gap between biophysically detailed muscle models and computational efficiency, JiSuJi has significant potential for advancing physics-based simulation of neuromuscular control in virtual bodies.

## Acknowledgements

Villum Fonden (36004) to IA; Carlsberg (CF22-0632) and Novo Nordisk Foundation (NFF20OC0063964) to CPHE; NIH (R01NS099375 and R01NS084844) to CPHE and SJS. This work used Expanse at the San Diego Supercomputer Center (SDSC) through allocation CTS180004 from the Extreme Science and Engineering Discovery Environment (XSEDE) (82), which was supported by National Science Foundation grant number #1548562 to QX and XZ.

## Methods

### 1. JiSuJi model

#### 1.1 Isometric muscle fiber force model

In this section, we introduce the chemomechanical model for calculating the muscle fiber force (*f*) in isometric condition in optimal length, L_0_ (the length at which muscle fibers generate the maximum isometric force) (83). The physiological process is divided into four steps: (1) the change of the permeability of calcium channel membrane to calcium due to the arrival of neural spikes; (2) the change of free calcium concentration in sarcoplasm (SP); (3) the binding and unbinding process of calcium to troponin and (4) the force generation through cross bridge cycling. The model incorporates a term accounting for CDI within calcium release dynamics, which dynamically modulates calcium release based on ISI, as informed by experiment measurements (51,84). With this component, the model can accurately capture the influence of ISI on calcium release: closely-spaced spikes result in less calcium release, and the release gradually recovers with longer intervals.

The input of the isometric muscle fiber force model is *imp(t)*, where *imp(t)* has the format of *t*_*1*_, *t*_*2*_, *t*_*3*_ …, representing the time instants of the arrival of neural spikes.

***Step 1***. The permeability of calcium channel membrane at the arrival of each spike (*k*) is assumed to be a rectangular pulse with the amplitude, *R*, and width, *t*_*w*_. When there are more spikes, according to the time interval between the spikes, the amplitude of the later spike is calculated through two coefficients: *c*_*amp*_ and *c*_*recover*_.

*c*_*amp*_ is used to consider the fact that the calcium release from consecutive spikes would decrease due to small ISIs as observed in experiments (51). The expression of *c*_*amp*_ is proposed as:

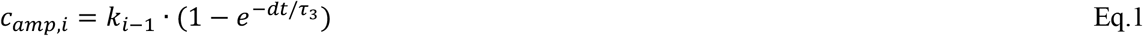

where *c*_*amp,i*_ is the coefficients for the *i* th spike; *k*_*i*−1_ is the permeability of calcium channel membrane at the arrival of the *i-1* th spike; *dt* is the time interval between *i* th and *i*-1 th spike; *τ*_3_ is the model constant. According to Eq.1, when *dt* =0, *c*_*amp,i*_ =0; when *dt* → ∞, *c*_*amp,i*_ = *k*_*i*−1_. The physical meaning is that when the two spikes are very close, the calcilum release resulted from the later spike would be close to zero. In the extreme condition that the later spike arrives at the same time with the previous spike, it will not result in additional calcium release. When the interval of the two spikes is large, the calcium release from the later spike is less affected.

*c*_*recover*_ is used to consider the fact that the calcium release will recover with time (84). A constant recover rate is assumed:

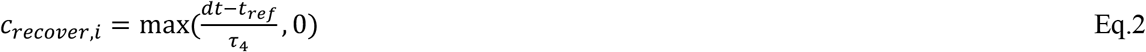

where *τ*_4_ is model constant; *t*_*ref*_ is the refractory period, which is fixed to be 0.82ms according to the measurement in the syringeal muscle (33). Eq.2 indicates a linear recover of calcium release with *dt*. The recovery would be zero if the time interval between the two spikes is smaller than the refractory period.

Therefore, *k*_*i*_ resulted by the *i* th spike arriving at *t*_*i*_ has the expression:

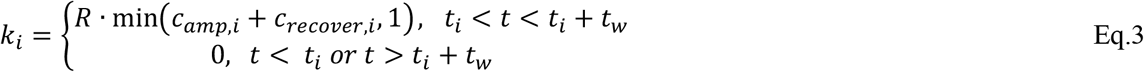

The expression of *k* due to multiple spikes is simply the summation of each *k*_*i*_:

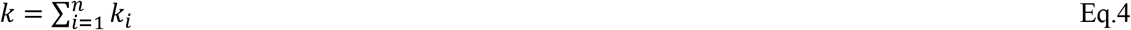

***Step 2***. The following Steps 2-4 follow the method in Wexler el al. (40). The concentration of SP calcium is affected by calcium release and uptake by SR and the binding and unbinding from the troponin:

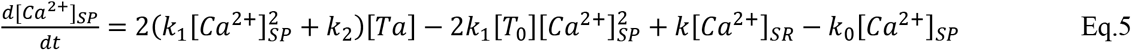

where [*Ca*^2+^]_*SP*_ is the free calcium concentration in sarcoplasm. The first two terms on the right hand side (RHS) represent the calcium binding and unbinding process from the troponin: 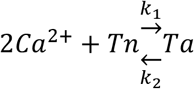, where *k*_*1*_ and *k*_*2*_ represent forward and backward reaction rate for calcium binding reaction, respectively; [*Ta*] is the concentration of activated troponin, which are in the *Ca*^2+^-troponin complex; [*Tn*] is troponin available for the binding process; [*Tn*]+[*Ta*] equals to the total concentration of troponin [*T*_*0*_], which is a model constant, fixed to be 140×10^−6^ M (24). The third and fourth term in Eq.5 represent the release and uptake of calcium from SR, respectively. [*Ca*^2+^]_*SR*_ is a model constant representing the concentration of free calcium in SR, which is hardly depleted. We set [*Ca*^2+^]_*SR*_ to be 0.001M (50,85). *k*_*0*_ is uptake rate of calcium by SR.

***Step 3*** Binding and unbinding of calcium to troponin:

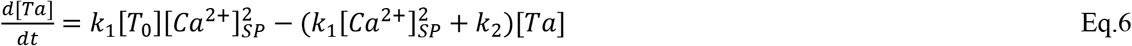

This is the same process described by the first two terms on the RHS of Eq.5.

***Step 4*** Force mechanics:

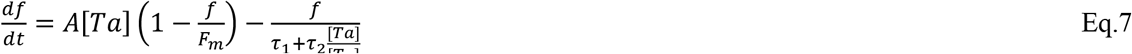

where *τ*_1_ and *τ*_2_ are time constants related to force decay; *F*_*m*_ is a model constant related to muscle force; *A* is a model constant of the force increase; *f* is the force response of single fiber to stimulation in isometric condition and optimal length. In Eq.7, the force generation is described as a serial connected spring-damper-motor system and the contractile speed of the muscle is assumed to be positively related to [*Ta*]. A detailed description can be refered to Wexler et al (40).

**Table S1** summarizes the eleven model parameters. They were obtained through a generic algorithm based optimization method by minimizing model predictions and experiment measurements (86). We benchmarked our model against two experimental dataset that measures the isometric force generated by isolated muscle bundles: a previously published isometric force dataset measuring the zebra finch DTB muscle response to twenty-five three-spike patterns (33); and a new experimental dataset measuring the Bengalese finch VS muscle response to ten short duration stimulations with varied ISI and long duration stimulation with constant ISIs.

For the DTB muscle of the zebra finch, the force response of five three-spike patterns and three long duration stimulations were used for optimization. The three-spike patterns included (1) [0, 1, 10] ms; (2) [0, 9, 10] ms; (3) [0, 2, 5] ms; (4) [0, 1.66, 3.3] ms; (5) [0, 1.25, 2.5] ms. The numbers in the square brackets represent the timing of the neural spikes. The ex vivo force response data were adopted from (33). The long duration stimulations were 400, 600, 700Hz with constant ISIs. Since no long duration stimulation measurements were available for the same muscle, we adopted the force responses of the long duration stimulation on the DTB muscle of a different zebra finch reported in (33). To make the force amplitudes compatible, we scaled the long-duration force responses by matching the force output of the first three spikes to the force responses of the corresponding three-spike patterns measured in our current muscle at the same frequencies with constant ISIs. For the VS muscle of the Bengalese finches, the transient force response of three two-spike patterns and two long duration stimulations were used for optimization. The two-spike patterns include (1) [0, 2] ms; (2) [0, 8] ms; (3) [0, 10] ms, with the two numbers in the square brackets represent the timing of the neural spikes. The long duration stimulations were 400 and 500Hz simulations with constant ISIs. We sampled the data in each case in a frequency of 100kHz. In each sampling point, the difference between the predicted force and measured force was calculated. The objective function was calculated as the sum of the absolute difference, with larger weight put on the time duration when the force response was near the peak value. The obtained model parameters are listed in **Table S1**.

#### 1.2 JiSuJi – A finite element framework

JiSuJi – a finite element framework was developed by intergrating the current muscle fiber force model with our previous 3D finite element framework (46,47). The new model calculated the passive and active tissue stress on 3D finite element models. We neglect the spatial variation of the muscle fibers, and assume that the muscles fibers are evenly distributed and recruited, and all the neurons are stimulated synchronously with external electrical stimuli to mimick the experimental setup. The total stress generated in the muscle is the sum of the passive component (*σ*_*pas*_) and the active component (*σ*_*act*_).

##### Passive component

The passive component employs a fiber-reinforced material (87), with parameters optimized to match experimentally measured passive stress curves (**Fig. S2**). The strain energy function of the passive material was defined as:

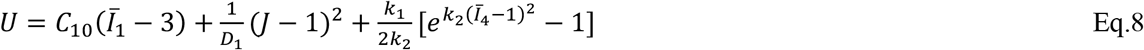

where *U* is the energy function; 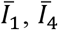 are the reduced invariants of the reduced Cauchy-Green tensor, respectively; *D*_1_ is material constant related to compressibility; *J* is the Jacobian determinant of the deformation; *C*_10_ is the material constant for the neo-Hooke term; *k*_1_, *k*_2_ are material constants for the exponential term (87).

The passive stress (*σ*_*pas*_), represented as the second Piola-Kirchhoff stress tensor *S*_*KL*_, is calculated as (87,88)

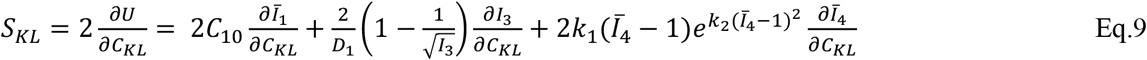

where *C*_*KL*_ is the right Cauchy-Green tensor; *I*_3_ is the third invariant of *C*_*KL*_.

To optimize model constants *C*_10_, *k*_1_ and *k*_2_, static finite element simulations were conducted on a rectangular bar at four different strain levels, as illustrated in **Fig. S2a**. A genetic algorithm–based optimization tool was employed to identify the set of model constants that minimized the difference between the simulated and measured passive stress values. The optimized *C*_10_, *k*_1_ and *k*_2_ were listed in **Table S3**.

##### Active component

The calculation of the active stress (*σ*_*act*_) follows the common assumption that the force–length and force– velocity relationships are independent of isometric muscle force production (20,89) and are modeled separately. We adopted previously developed models for both relationships(46,47), and their parameters were optimized individually to match experimental measurements (**Fig. S2bc**).

The total active stress is given by

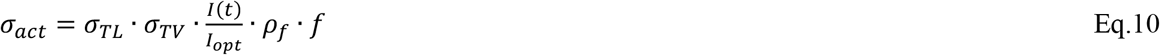

where *σ*_*TL*_, *σ*_*TV*_ are coefficients of length and velocity dependency, respectively; *I*_*opt*_ is the optimal current/voltage intensity, which is the lowest current intensity that all fibers can be recruited; *ρ*_*f*_ is the fiber density, equaling to 1600 fibers/mm^2^ for Bengalese finch syringeal muscle (90) and 1060 fibers/mm^2^ for zebra finch syringeal muscle (36); *f* is the isometric force response of a single muscle fiber to the input stimulation at optimal length, calculated in Eq.7.

The length dependency coefficient (*σ*_*TL*_) of the active stress is caluclated as Eq.11 (69):

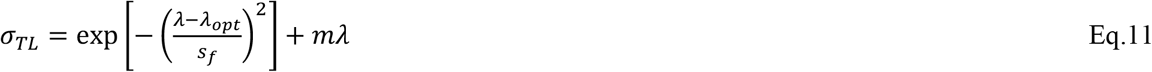

where *λ* is the stretch calculated as the current muscle length divided by the initial length (*L*_0_); *λ*_*opt*_ is the optimal stretch, reported to be 1 according to (48); *s*_*f*_ and *m* are model constants that were optimized to fit the experimentally measured active stress–length relationship reported in (48). The optimized values of *s*_*f*_ and m are listed in **Table S3** and the resulting *σ*_*TL*_ –λ curve is shown in **Fig. S2b**.

The velocity dependency coefficient (*σ*_*TV*_) of the active stress is caluclated as Eq.12 (69):

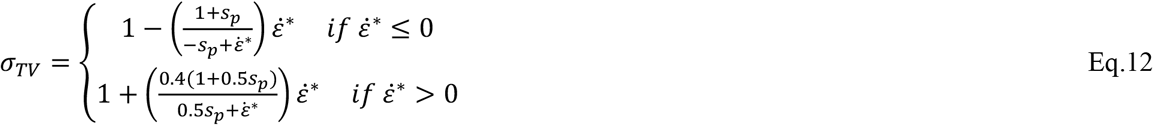

where *s*_*m*_ is model constant; 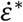 is the normalized strain rate calculated as v/v_max_, where v is the shortening velocity of the tissue and v_max_ is the maxmium shortening velocity, which was reported to be 33.6 L_0_/s from the measurement **(Method 2.4)**.

The velocity – stress relationship was obtained through isovelocity test by shortening the tissue from 1.1L_0_ to L_0_ at various velocities from 0.25 L_0_/s to 20L_0_/s. The active stress corresponding to the time instant when the muscle was back to the original length was recorded (**Method 2.4**). By normalizing this value using the active stress right before the shortening period, 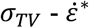 relationship was obtained from the measurement. The model constant *s*_*m*_ was optimized to fit the obtained 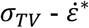 relationship. The optimized *s*_*m*_ is listed in **Table S3** and the resulting *v/v*_*max*_ – *σ*_*TV*_ curve is shown in **Fig. S2c**.

To capture the relaxation in active stress following a sudden change in velocity, a relaxation model employing two coupled ordinary differential equations were applied to *σ*_*TL*_ · *σ*_*TV*_ (69):

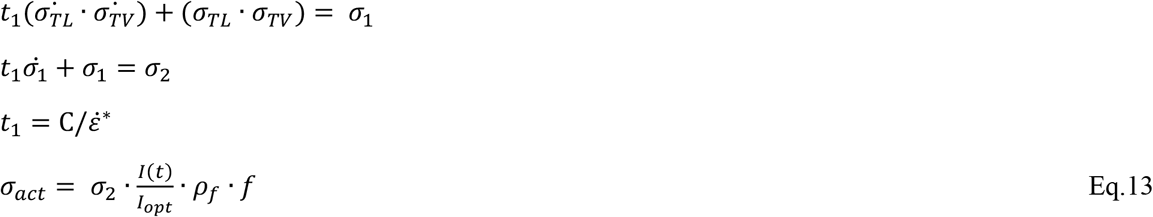

where *σ*_1_ is the intermediate response; *σ*_2_ is the relaxed active stress coefficient. *σ*_*act*_ is the final active stress with relaxation; *t*_1_ is relaxation time constant; C is a model constant and was optimed to fit the measured active stress reposne during shortening period for two shortening velocities of 5L_0_/s and 20L_0_/s (**Fig. S2d**). The optimzied value of C is listed in **Table S3**.

### 2. Experimental data

#### 2.1 Isometric force measurement on the zebra finch syrinx muscle

Force responses of twenty-five three-spike stimulation patterns were measured as described in detail in (33) on the dorsal tracheobronchial muscle (DTB) in male zebra finches. Briefly, the three-spike stimulation patterns were based on stimulation frequencies of 200, 400, 600, and 800Hz, with the timing of the middle spike moved from the centered position toward the first or the third (last) spike. For JiSuJi implementatiom, we assumed the fiber density was 1060 fibers/mm^2^ based on ref. (36).

#### 2.2 In vivo [Ca2+] signal transduction dynamics

We previously quantified [Ca^2+^] in zebra finch syringeal muscle (vental tracheobronchial muscle VTB)as a function of temperature (55). In brief, force development curves of single and multiple stimuli (100 ms, 100 Hz trains) were measured at a range of temperatures (10–39°C) at optimal stimulus amplitude and muscle length. Next, we inhibited actomyosin interaction to avoid movement artifacts for calcium measurements using BTS (91). To image intracellular calcium, we used Mag-Fluo-4 as high-affinity calcium indicator (92) and measered light with a photomultiplier (PTM) mounted on an inverted microscope (Nikon eTI, DFA, Glostrup, Denmark). Force, stimulus and PTM signals were sampled at 60 kHz (National Instruments USB6259) and analyzed in Matlab.

The PTM signal was highpass filtered (30 Hz, 2^nd^ order butterworth filter, filtfilt implementation) to remove slow changes in the PTM signal. DFF was defined as the PTM signal minus the mean of 100 samples prior to stimulation. To calculate FWHM of the DFF signals we also applied a lowpass filter (500 Hz, 2^nd^ order butterworth filter, filtfilt implementation) and detected the DFF peaks after stimulation pulses.

#### 2.3. Isometric force measurement on the Bengalese finch VS muscle

Force responses to both short (one to three spikes) and long duration stimulation patterns (100 to 500Hz) were measured on dissected muscle fiber bundles of ventral syringeal muscle (VS) in male Bengalse finches. In the short duration stimulation pattern, the spike intervals were in the range of 2 to 10 ms.

#### 2.4 Isovelocity test

A detailed description of the isovelocity test has been provided in ref. (48). In brief, the DTB muscle bundle was dissected out of an extracted zebra finch syrinx. The muscle bundle was mounted on a force transducer (400A, Aurora Scientific, Aurora, ON, Canada) at one end and a high-speed length controller (322C, Aurora Scientific) at the other to enable the measurement of force and dynamic control of muscle length. The muscle was initially stretched to 1.1*L*_*0*_ and tetanized to elicit maximal isometric force using a stimulation frequency of 600Hz. During stimulation, the muscle was shortened back to *L*_*0*_ at velocities ranging from 0.25 to 20 L_0_/s using the high-speed length controller. Stimulation ceased once the tissue returned to *L*_*0*_. A parallel run at each velocity was conducted without stimulation to isolate passive force. Active muscle force at each velocity was obtained by subtracting passive force from total force.

#### 2.5. 3D rendering of a syringeal muscle preparation

To obtain the accurate geometry of the syringeal muscle fiber bundle, one DTB preparation was 3D scanned following the experimental procedure. The preparation was set to approximately *L*_*0*_ in a petri-dish. This preparation was then scanned using an ultrasound imaging system (Vevo F2, Fujifilm, Tokyo, Japan). Scans were made along both the width and length of the muscle bundle at the highest in-built resolution setting (20 µm). The images were then imported into Mimics (Materialise, Belgium) to reconstruct a 3D representation of the muscle bundle.

### 3. Simulation – Experimetal comparison metrics

The coefficient of determination (*R*^2^) was calculated as 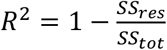, where the residual sum of squares was calculated as *SP*_*res*_ = ∑_*i*_(*y*_*i*_ − *f*_*i*_)^2^ = ∑_*i*_ *e*_*i*_^2^, where *y*_*i*_ is the observed data. *f*_*i*_ is the corresponding fitted data. The total sum of squares was calculated as 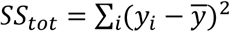, where 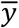 is the mean of the observed data.

#### Fusion index and fusion frequency

Fusion index (FI) was calculated for each stimulation frequency using the method in (56). The index represents the ratio of the minimum to the maximum force produced between two consecutive stimuli. A FI value of 1 indicates complete tetanus, where all individual twitches have merged into a single sustained contraction. Lower fusion index values indicate less fusion, suggesting that individual twitches remain distinct. Fusion frequency (f_fused_) corresponds to a FI of 0.9.

#### Steady force and t_90_

Steady force is defined as the time-averaged muscle force during the late phase of long duration stimulation, when the cycle-averaged force no longer changes significantly over time. For the force response of the DTB muscle of the zebra finch, the steady force was calculated using the data from 15ms to 25ms in Fig. 1c. Regarding the force response of the VS muscle of the Bengalese finch, for the Wilson model, the steady force was computed as the mean force between 150 ms and 180 ms, while for all other cases, it was calculated over the 60 ms to 90 ms window. t_90_ is defined as the time point at which the muscle force reaches 90% of the steady force. Since muscle force can oscillate at lower stimulation frequencies, a cycle-averaged force was computed for each cycle. A smooth force-time curve was then generated by fitting the cycle-averaged data using a B-spline in MATLAB. The value of t_90_ was determined based on this smoothed curve.

## Notes

### Competing Interest Statement

The authors have declared no competing interest.

