## Supplemental material for "JiSuJi, a virtual muscle for small animal simulations, accurately predicts force from naturalistic spike trains"

### SUPPLEMENT MATERIAL

Weili Jiang[1], Iris Adam[2], Nicholas W. Gladman[2,4], Sam Sober[3], Qian Xue[1], Coen P.H. Elemans[2]*, Xudong Zheng[1]*

[1]Mechanical Engineering Department, Rochester Institute of Technology, Rochester, NY, United States

[2]Sound Communication and Behavior Group, Department of Biology, University of Southern Denmark, Denmark

[3]Department of Biology, Emory University, Atlanta, GA, United States

[4]Present address: Department of Musculoskeletal and Ageing Science, Institute of Life Course and Medical Sciences, University of Liverpool, Liverpool, United Kingdom

**Supplementary Text**

In this section we described the Wilson model (1) and the Wexler model (2), accroding to which the force responses were caclulated (**Fig 2** in the main text).

**1. Models description**

A detailed description of the Wilson model was in (1). Here we listed the core equations. The model takes a pulse train u(t) as input where:

$$u(t) = \sum_{i=1}^{n} \delta(t - t_i) \quad \text{Eq.S1}$$

where n is the number of input pulses, $t_i$ the time the *i*th pulse occurs.

$$\dot{C}_N(t) + \frac{C_N(t)}{\tau_c} = u(t) \quad \text{Eq.S2}$$

$$x(t) = \frac{C_N(t)^m}{C_N(t)^m + k^m} \quad \text{Eq.S3}$$

$$\dot{F}(t) + \frac{F(t)}{\tau_1 + \tau_2 x(t)} = Ax(t) \quad \text{Eq.S4}$$

The model has six parameters: $\tau_c$, $\tau_1$, $\tau_2$, m, k and A.

A detailed description of the Wexler model was in (2). Here we listed the core equations. The model input is k, which is either 0 or 1. It is 1 for 4ms following the arrival of each action potenital. The response of $[Ca^{2+}]_{SP}$, [Ta] and $f$ are described using Eq.S5 – 7.

$$\frac{d[Ca^{2+}]_{SP}}{dt} = 2(k_1[Ca^{2+}]_{SP}^2 + k_2)[Ta] - 2k_1[T_0][Ca^{2+}]_{SP}^2 + k[Ca^{2+}]_{SR} - k_0[Ca^{2+}]_{SP} \quad \text{Eq.S5}$$

$$\frac{d[Ta]}{dt} = k_1[T_0][Ca^{2+}]_{SP}^2 - (k_1[Ca^{2+}]_{SP}^2 + k_2)[Ta] \quad \text{Eq.S6}$$

$$\frac{df}{dt} = A[Ta]\left(1 - \frac{f}{F_m}\right) - \frac{f}{\tau_1 + \tau_2\frac{[Ta]}{[T_0]}} \quad \text{Eq.S7}$$

The value of $\tau_1$ was found by performing linear regression for the force decay after the tetanus as suggested in (2). Thus the model has seven parameters - $k_1$, $k_2$, $k_0$, $[T_0]$, A, $F_m$, $\tau_1$, $\tau_2$ – that the values need to be obtained from the optimization process.

### 2. Model parameters and performance

To obtained the model constants in the Wilson model and the Wexler model. The same dataset from the measurement of the Bengalese VS muscle, the same genetic algorithem optimization method, and the same objective function with the ones used for JiSuJi were employed. **Table S3** lists the model parameters obtained from the optimization process. **Figure S1** shows the responses of the models to the short duration stimulation with varied ISIs. The models responses to the long duration stimulation with constant ISIs and the comparison with the measurments are shown in **Figure 2** in the main text.

### Supplementary Tables

**Table S1. Parameters of isometric force model.** The optimized values for the dorsal tracheobronchial (DTB) muscle from the zebra finch and the ventral syrinx (VS) muscle from Bengalese finch were listed.

| Variable | Unit | Description | Value | |
|---|---|---|---|---|
| | | | DTB muscle Zebre finch | VS muscle Bengalese finch |
| $\tau_1$ | ms | Model constant related to force relaxation | 1.97 | 2.75 |
| $\tau_2$ | ms | Model constant related to force relaxation | $7.68\times10^4$ | $4.0\times10^3$ |
| $\tau_3$ | ms | Model constant related to the decreased calcium release from consecutive spikes | 3.84 | 1.08 |
| $\tau_4$ | ms | Model constant related to the recover of calcium release | 0.91 | 8.33 |
| $k_1$ | $M^{-2}\cdot ms^{-1}$ | Forward reaction rate of calcium binding reaction | $3.31\times10^8$ | $7.71\times10^8$ |
| $k_2$ | $ms^{-1}$ | Backward reaction rate of calcium binding reaction | 0.69 | 1.51 |
| $k_0$ | $ms^{-1}$ | Uptake rate of calcium by SR | 0.46 | 0.28 |
| $A$ | $mN\cdot ms^{-1}\cdot M^{-1}$ | Model constant related to force increase | $9.97\times10^4$ | $4.4\times10^3$ |
| $F_m$ | mN | Model constant related to muscle force | 0.007 | 0.038 |
| $t_w$ | ms | Width coefficient of *k* resulted from each spike | 1.16 | 1.34 |
| $R$ | $ms^{-1}$ | Amplitude coefficient of *k* resulted from each spike | $5\times10^{-4}$ | $1.8\times10^{-3}$ |

**Table S2. Model parameters of the Wexler model and the Wilson model**. The detailed models were described in (2) and (1), respectively.

| Wexler model | | | | | | | |
|---|---|---|---|---|---|---|---|
| | $k_1$ | $k_2$ | $k_0$ | A | $F_m$ | $T_0$ | $\tau_2$ |
| Value | $8.1\times10^{8}$ | 1.7 | 0.6 | $4.5\times10^{6}$ | 39.5 | $1.9\times10^{-4}$ | $7.1\times10^{3}$ |

| Wilson model | | | | | | |
|---|---|---|---|---|---|---|
| | $\tau_c$ | $\tau_1$ | $\tau_2$ | m | k | A |
| Value | $3.7\times10^{-3}$ | $5.5\times10^{-3}$ | $2.2\times10^{-2}$ | 9.5 | 0.76 | $1.48\times10^{3}$ |

**Table S3. Model parameters for the passive and active material properties.** $\lambda_{opt}$ and $v_{max}$ were measured from experiment. The other parameters were obtained through optimization.

| | |
|---|---|
| Passive component (Eq.8) | |
| | $C_{10}$=2.92 $k_1$=0.877 $k_2$=19.482 |
| Active component (Eq.12 – 14) | |
| $\sigma_{TL}$ | $s_f$=0.329, $m$=0, $\lambda_{opt}$=1 |
| $\sigma_{TV}$ | $s_p$= 0.17, $v_{max}$=33.6$L_0/s$ |
| C | 0.56 |

**Supplementary Figure**

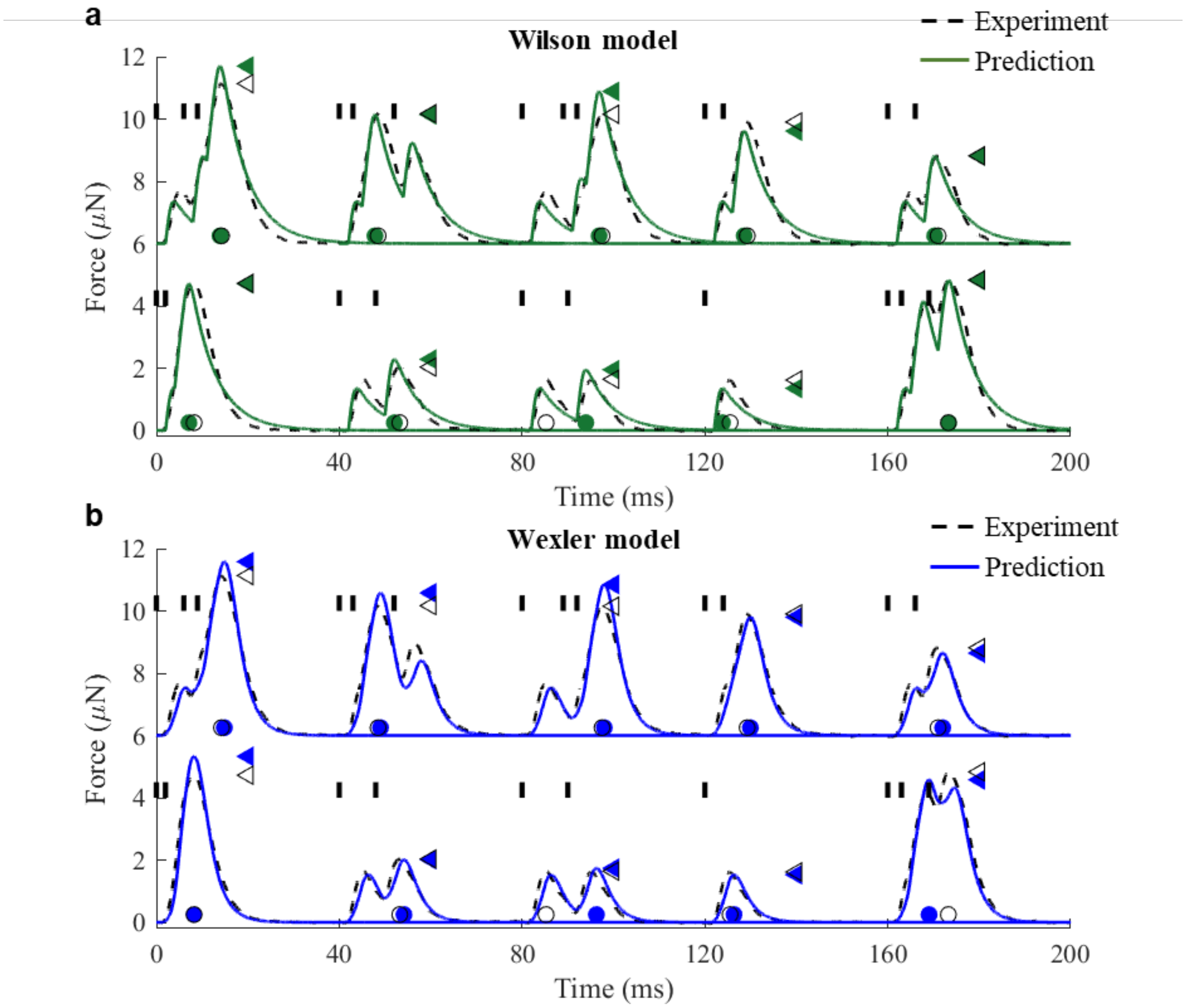


**Figure S1. Predicted forces by state-of-the-art models. a**, Performance of Wilson model and **b**, Wexler model compared to experimental data in response to ten short duration stimulation patterns (vertical short bars) in the Bengalese finch ventral syringeal (VS) muscle. Time instants (circles) reaching the maximum forces (triangles) were denoted.

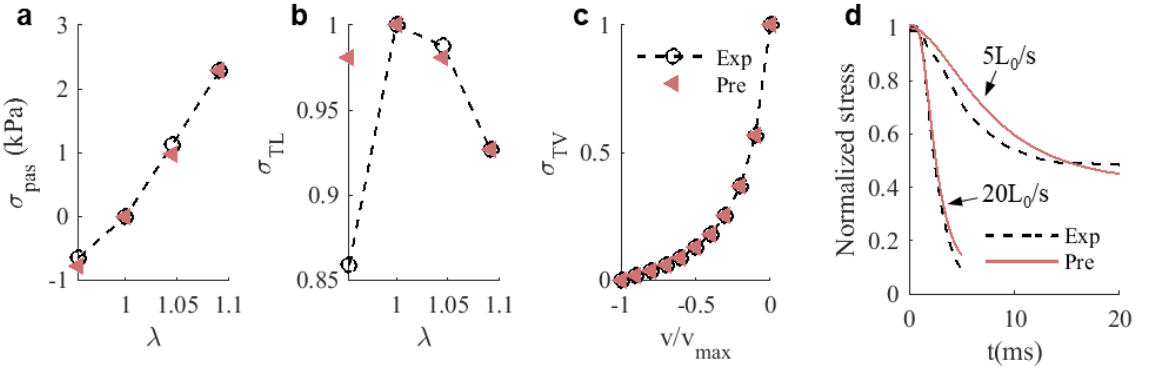


**Figure S2. Optimization of model parameters.** **a**, The passive stress – stretch ($\sigma_{pas}$ – λ) relationship, **b**, the active stress – stretch ($\sigma_{TL}$ – λ) relationship, **c**, the active stress – velocity ($\sigma_{TV}$ – $v/v_{max}$) relationship, and **d**, the tissue response in the shortening period at two shortening velocities.
